# Biosensor imaging of G protein mediated Signaling Pathways Activated Immediately Downstream of CaSR

**DOI:** 10.1101/2024.05.30.596581

**Authors:** Sergei Chavez-Abiega, Frank J. Bruggeman, J. Goedhart

## Abstract

Genetically encoded biosensors allow us to study events in single cells in real time. Here, we used biosensors with a read-out based on Förster resonance energy transfer (FRET) or intensity to investigate the main signaling pathways activated immediately downstream of the calcium sensing receptor (CaSR) in HEK-293 cells. CaSR responds to its endogenous ligand, calcium, to activate Gq and Gi protein dependent pathways. We demonstrate that CaSR activates Gq to promote intracellular calcium mobilization, and we monitored these two events simultaneously in the same cells. We also observed that CaSR activates the 3 Gαi subtypes and decreases cAMP levels in a Gi-dependent manner. Moreover, the use of negative allosteric modulator NPS-2143 can inhibit these signaling events and also rapidly disrupts them if added after receptor activation by calcium stimulation. In addition, the increases in calcium and cAMP levels were respectively enhanced when activation of Gi or Gq proteins was prevented. Finally, we provide evidence that CaSR does not couple to G13 proteins, and that activation of RhoA by CaSR is solely dependent on Gq/G11 activity.

## INTRODUCTION

The calcium-sensing receptor (CaSR) is ubiquitously expressed in the body, and mostly in kidneys and parathyroid glands, where it plays a fundamental role in calcium homeostasis. CaSR senses and keeps the levels of free circulating extracellular calcium at 1.1-1.3 mM. It does so by regulating the secretion of the parathyroid hormone and the reabsorption of calcium by the kidneys [1]. CaSR is an important clinical target for the treatment of secondary hyperparathyroidism, with two FDA-approved drugs in the market, cinacalet and etelcalcitide.

CaSR belongs to the class C of G protein-coupled receptors (GPCRs), characteristic for being obligate dimers and containing a large extracellular domain (ECD). CaSR is present in the membrane as a homo-dimer, but it is suggested that it can form heterodimers with other class C GPCRs [2,3]. The ECD is composed of two lobe-shaped domains, LB1 and LB2, that form the so-called Venus Fly Trap domain (VFT). The VFT domains allocate the binding sites for orthosteric ligands such as calcium, divalent and trivalent cations, poly-aminoacids, among others [4]. Upon ligand binding, the VFTs close, and the LB2 and CR domains of both protomers approach to each other and interact to form a large interface. As a result, the distance between the C-termini of the ECDs reduces from 83 Å to 23 Å, and it is proposed that this causes the rearrangement of the TMD that finally leads to coupling to downstream effectors [5]. The ECD of CaSR contains several sites for N-linked glycosylation, necessary for full maturation and expression in the membrane. The non-glycosylated monomer is 120 kDa, and the fully glycosylated mature monomer 160 kDa.

The CaSR couples predominantly to the G_q_ and G_i_ heterotrimeric G protein families [6], as reported by multiple studies, and some studies report coupling to G_13_ and G_s_ in specific cell lines. Previous studies in HEK-293 and bovine parathyroid cells have shown that activation of CaSR results in decrease of cAMP levels that is sensitive to Pertussis toxin and in binding of GTP to G_αi_ proteins. Here, we use live cell imaging with genetically encoded biosensors to examine at the single cell level which heterotrimeric G-proteins are activated by CaSR.

## RESULTS

### Gq and Gi activation by CaSR

We used fluorescence-based biosensors to study the activation of heterotrimeric G proteins and second messengers. First, we used FRET-based biosensors to quantify heterotrimeric dissociation of the Gα_q_, Gα_i1_, Gα_i2_ and Gα_i3_, from the βγ subunits [7,8] in HEK-293 cells stably expressing CaSR (HEK-CaSR). These biosensors show a FRET decreases upon activation, presumably through dissociation of the activated heterotrimer, as shown in Figure 1A. Using the Gq sensor, we observed decreases in the FRET ratio upon stimulation with increasing concentrations of calcium, as shown in Figure 1B-C. This effect is reversed upon the addition of the CaSR-specific negative allosteric modulator NPS-2143, and is specific to Gq activity, as shown by the preincubation with the Gq inhibitor YM-254890 [9].

**Figure 1.**
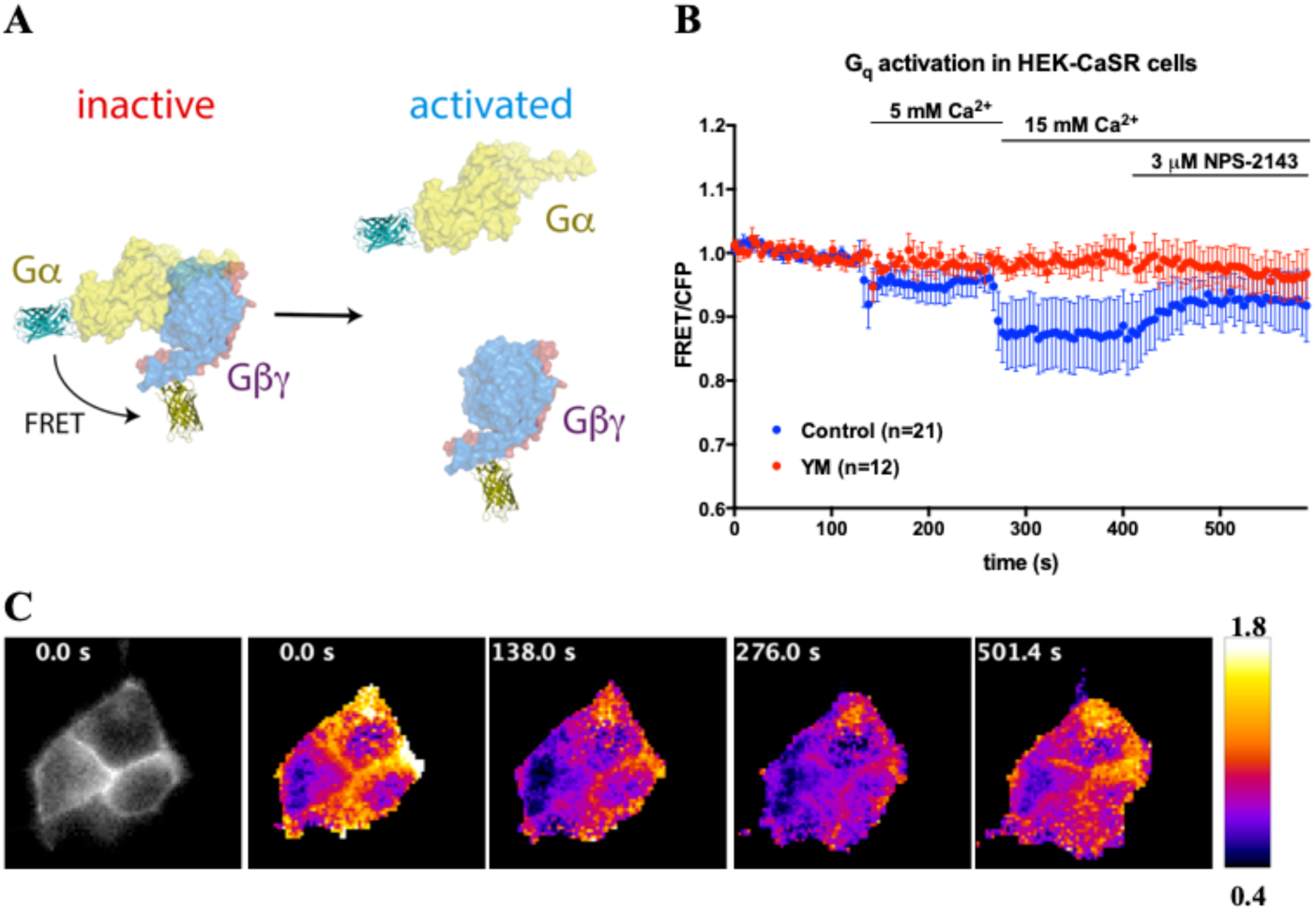
Activation of Gq by CaSR. A: Representation of the inactive and active states of the G protein biosensors. B: Average FRET/CFP ratios plotted over time. C: Not normalized FRET/CFP ratios from three cells at 4 different time points. Cells were stimulated with Ca2+ at the showed time points, and the response antagonized by NPS-2143. Cells were preincubated either with 1 μM YM-254890 or DMSO for 2 h prior to imaging. Time traces show the average ratio change of FRET/CFP fluorescence, and error bars show the 95% CI of the mean. Number of cells is specified for each condition.

For the Gq sensor, mNG has been shown to be the ideal acceptor for ratiometric imaging as it shows the highest dynamic range [7]. Given the similar biosensor architecture, we replaced cpVenus by mNG from the published Gi3 biosensor and compared the dynamic range using both acceptors in response to activation of the Gi-coupled designed receptor hM4D by the pharmacologically inert CNO in HEK-293 cells [10]. Surprisingly, the originally published biosensor showed significantly higher dynamic range than the mNG-variant (average FRET change of 0.20 vs 0.14), as shown in Figure 2. Therefore, we decided to use the originally reported Gi biosensors [8] for the remainder of this study.

**Figure 2.**
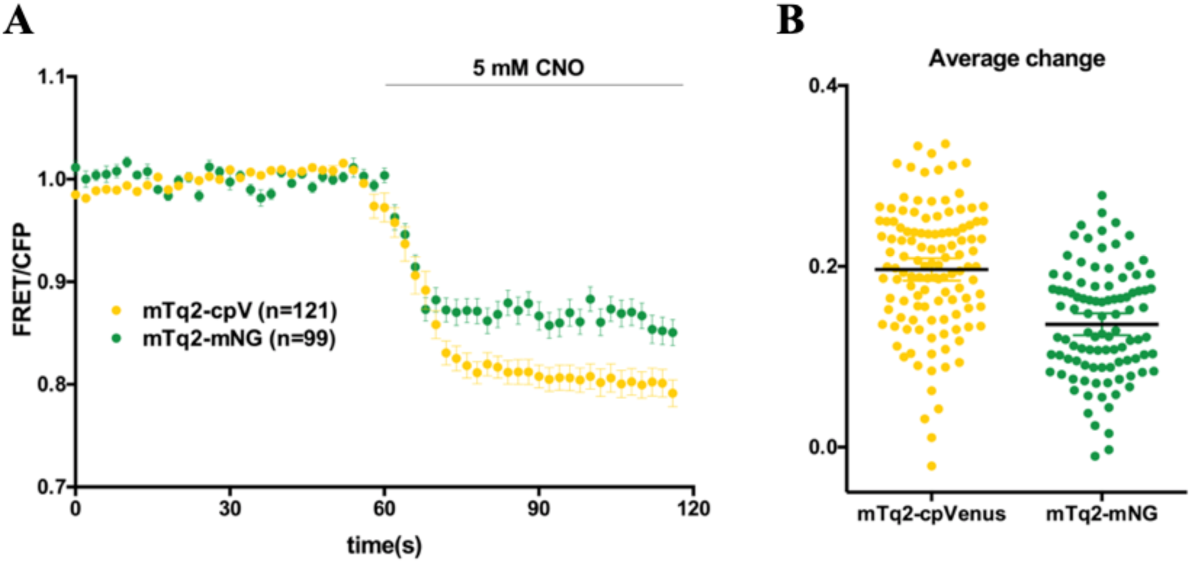
Comparison of cpVenus and mNeonGreen as acceptors for mTq2 in Gi3 sensor. A: Average FRET/CFP ratios plotted over time. B: Average ratio change per cell between 76 and 118 s after imaging started. Cells were stimulated with 5 mM CNO at the showed time point. Time traces show the average ratio change of FRET/CFP fluorescence, and error bars show the 95% CI of the mean. Number of cells is specified for each condition.

Ca^2+^ stimulation of HEK-CaSR cells transiently transfected with the FRET biosensors for the 3 Gαi subtypes resulted in comparable concentration-dependent FRET/CFP ratio decreases. This was reversed upon addition of NPS-2143 and prevented by preincubation with Gi inhibitor PTx, as shown in Figure 3.

**Figure 3.**
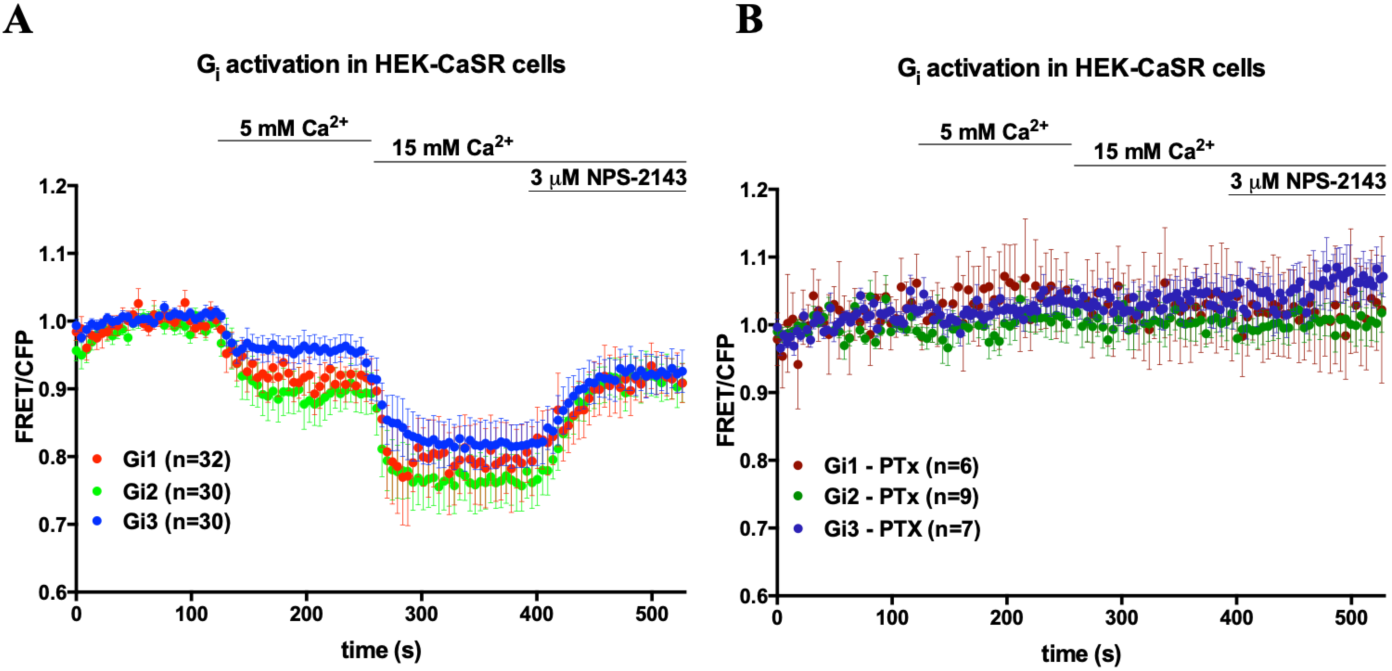
Activation of Gi by CaSR. A: Average FRET/CFP ratios plotted over time. B: Average FRET/CFP ratios plotted over time in the presence of PTx. Cells were stimulated with Ca^2+^ at the showed time points, and the response antagonized by NPS-2143. Cells were preincubated with 100 ng/mL PTx overnight prior to imaging. Time traces show the average ratio change of FRET/CFP fluorescence, and error bars show the 95% CI of the mean. Number of cells is specified for each condition.

### Calcium signaling by CaSR

Next, we studied several representative downstream effectors of these signaling pathways. The most relevant Gα_q_ signaling pathway involves the activation of PLCβ, which results in increased levels of DAG and Ca^2+^_i_. We decided to look into changes in Ca^2+^_i_ upon CaSR activation using the R-GECO1 biosensor [11]. This biosensor is based on a circularly-permutated mApple, and changes in Ca^2+^_i_ concentrations result in up to 16-fold changes in fluorescence intensity. Stimulation with 10 mM calcium results predominantly in rapid transient responses, as shown in Figure 4A. However, there are noticeable differences in the kinetics of the responses. The signals in most cells decrease and reach a plateau within 3-4 min after stimulation. Some cells show oscillations or intensity changes prior to stimulation, as observed in Figure 4E.

**Figure 4.**
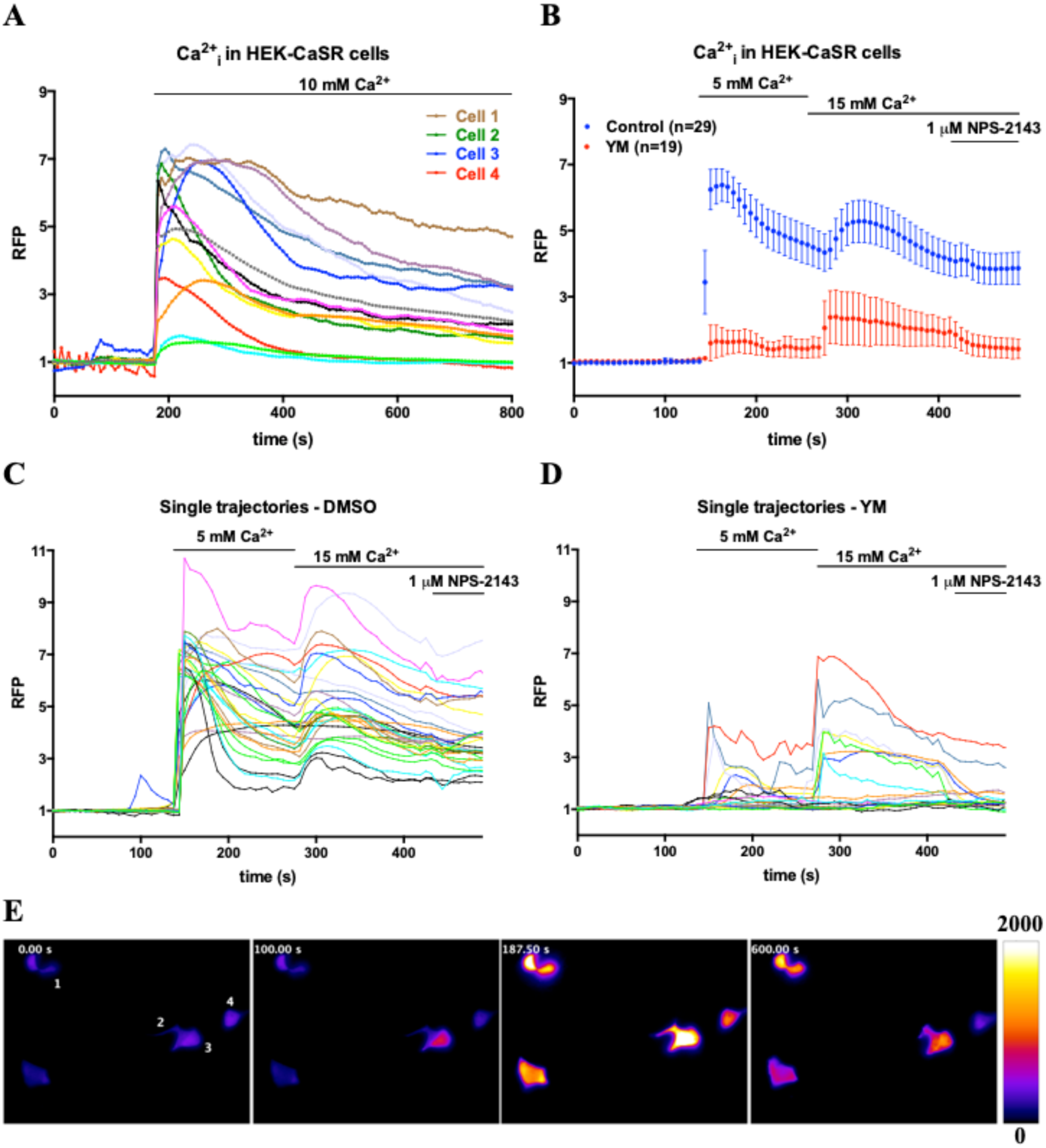
Calcium signaling by CaSR in HEK-CaSR cells transfected with R-GECO1. A, E: Cells were stimulated with 10 mM Ca2+. A: Single-cell normalized RFP intensity trajectories. E: RFP intensity counts at 4 different time points. B-D: Cells were stimulated with 5 and 15 mM Ca2+, and 1 μM NPS-2143. B: Average RFP intensity normalized to baseline in the presence of DMSO or YM. C-D: Single-cell trajectories for both conditions. Cells were stimulated at the showed time points with Ca2+, and the responses antagonized by NPS-2143. Cells were preincubated 2h prior to imaging with DMSO or YM. Time traces show the normalized RFP fluorescence intensity, and error bars show the 95% CI of the mean. Number of cells is specified for each condition.

When the cells were sequentially stimulated with 5 and 15 mM calcium, the magnitude of the second peak was similar or lower than the first one, as shown in Figure 4B. When Gq activity was inhibited, the majority of cells did not respond to calcium, and in those that responded, the second peak was higher. NPS-2143 had little or no effect in decreasing the signals.

We showed previously that Gq activity increases when calcium concentrations are applied consecutively, and it is stable for at least 2 minutes. Changes in intracellular calcium are, on the other hand, transient. Since the quantification of R-GECO1 requires only the RFP channel, we decided to image simultaneously Gq activity and Ca^2+^_i_ mobilization and in this manner follow two different events within the same signaling pathway in the same cell. Given that 5 mM calcium caused a higher response that the following addition of 10 mM calcium, we decided to sequentially stimulate the cells with 4, 8, 12, and 16 mM calcium. Upon stimulation with 4 and 8 mM calcium, the intracellular calcium levels increased and showed comparable amplitude, while subsequent calcium additions did not have a significant effect, as shown in Figure 5A. On the other hand, Gq activity increased to reach a maximum between 8 and 12 mM calcium, as observed in Figure 5B. Since the calcium mobilization responses showed such variability with the sequence of calcium concentrations used, we decided to increase the calcium concentration in smaller steps. Figure 5F shows calcium mobilization after adding 0.5 mM calcium, and the responses of concentrations up to 8 mM calcium were comparable, with 12 mM causing the highest peak.

**Figure 5.**
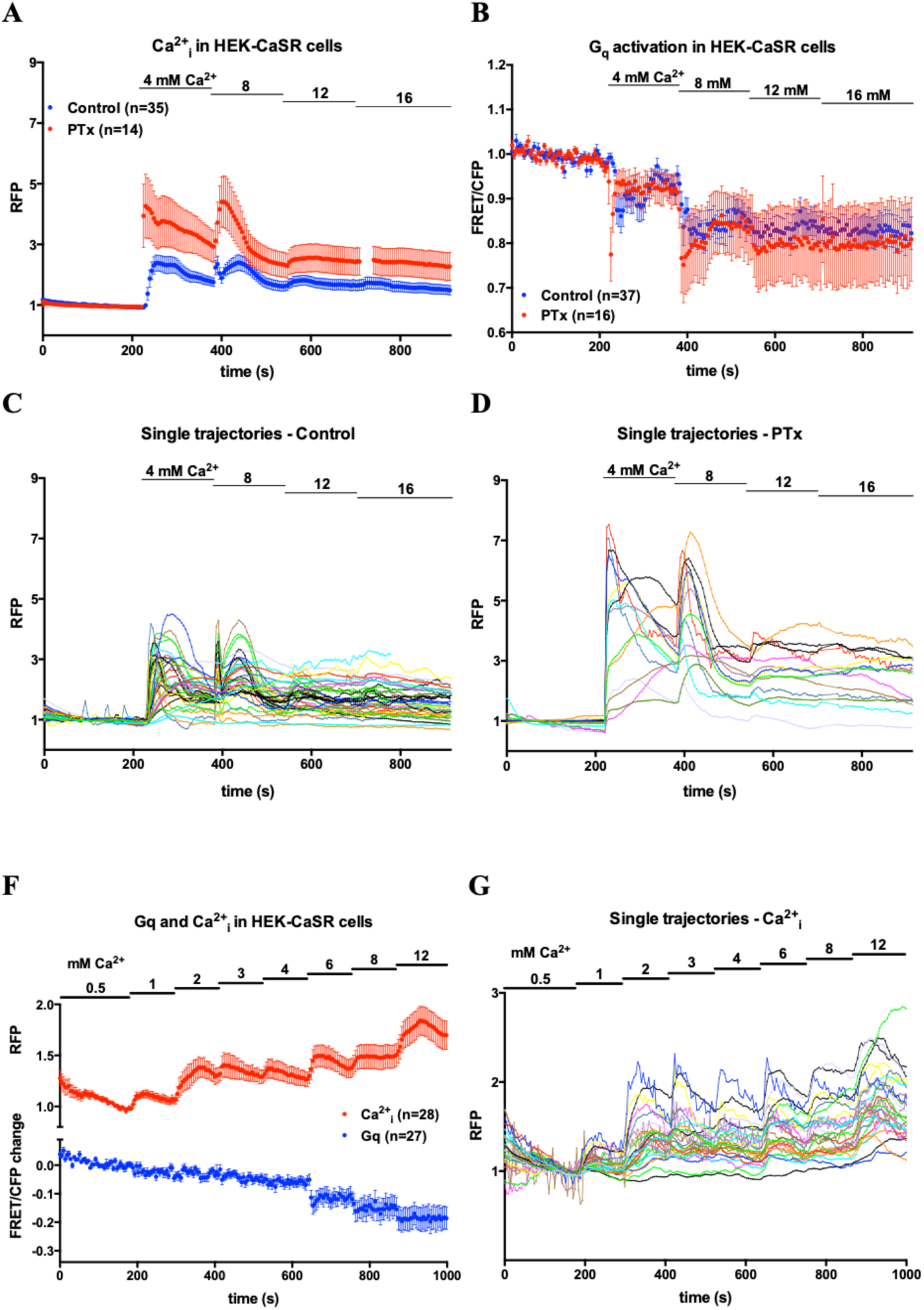
Calcium signaling by CaSR in HEK-CaSR cells transfected with R-GECO1 and Gq sensor. A-D: Cells were stimulated with various calcium concentrations in the presence and absence of PTx. A: R-GECO1. B: Gq sensor. C-D: Single-cell normalized RFP intensity trajectories. F-G: Cells were stimulated with various calcium concentrations. F: R-GECO1 and Gq. G: Single-cell R-GECO1 trajectories. Cells were stimulated at the showed time points with Ca2+. For PTx, cells were preincubated overnight with PTx. Time traces show the normalized RFP or FRET/CFP ratio, and error bars show the 95% CI of the mean. Number of cells is specified for each condition.

Since CaSR couples to both Gq and Gi proteins, we explored whether Gi inhibition could have an effect on the Gq activity or calcium mobilization. As shown in Figures 5B and 5D, Gi inhibition increased the amplitude of calcium mobilization responses, but it did not translate in clear increased coupling to Gq. Last, we explored the effect of even smaller sequential calcium increases up to 12 mM calcium. Under these conditions, shown in 5F-G, we observed a clear average increase in calcium mobilization between stimulation with 8 and 12 mM calcium.

### cAMP signaling by CaSR

G_i_ activity inhibits various types of adenylyl cyclases (ACs), resulting in decreased cytosolic cAMP levels and therefore less EPAC activity. Since we observed activation of Gi by the CaSR, we decided to look into cAMP levels. We used a biosensor that is based on EPAC (exchange factor directly activated by cAMP), showing high FRET in the absence of cAMP, and low FRET upon binding of cAMP. Calcium stimulation of HEK-293 cells transfected with CaSR (HEKwt-CaSR) and EPAC sensors H74 or H188 [12,13] did not result in any significant FRET change, with or without G_i_ inhibition, as shown in Figure 6A. We hypothesized that we did not observe any increase in FRET because the level of cAMP in resting cells is relatively low, and a further decrease by G_i_ activation could escape the detection limit of the sensor. We attempted to overcome this by using a high-affinity EPAC sensor, H187 [13], but the results were comparable to those with the other sensors (data not shown). We then decided to increase the cAMP level in the cells prior to calcium stimulation, by adding forskolin, which causes ubiquitous activation of ACs. As observed in Figure 6B, forskolin causes a very large decrease in FRET, as a result of increased cAMP. Stimulation with 15 mM calcium caused an increase in FRET with the H74 and H188 biosensors, as shown in Figure 6 B-C. This change was reversed by NPS-2143 and prevented by Gi inhibition with PTx. In HEK-CaSR cells transfected with H188, stimulation with 5 and 15 mM calcium resulted in a concentration dependent increase in FRET. Inhibition of Gq increased the measured FRET change upon calcium stimulation, as shown in Figure 6D, which resembles our observation of Gi inhibition enhancing the mobilization of Gq-dependent intracellular calcium.

**Figure 6.**
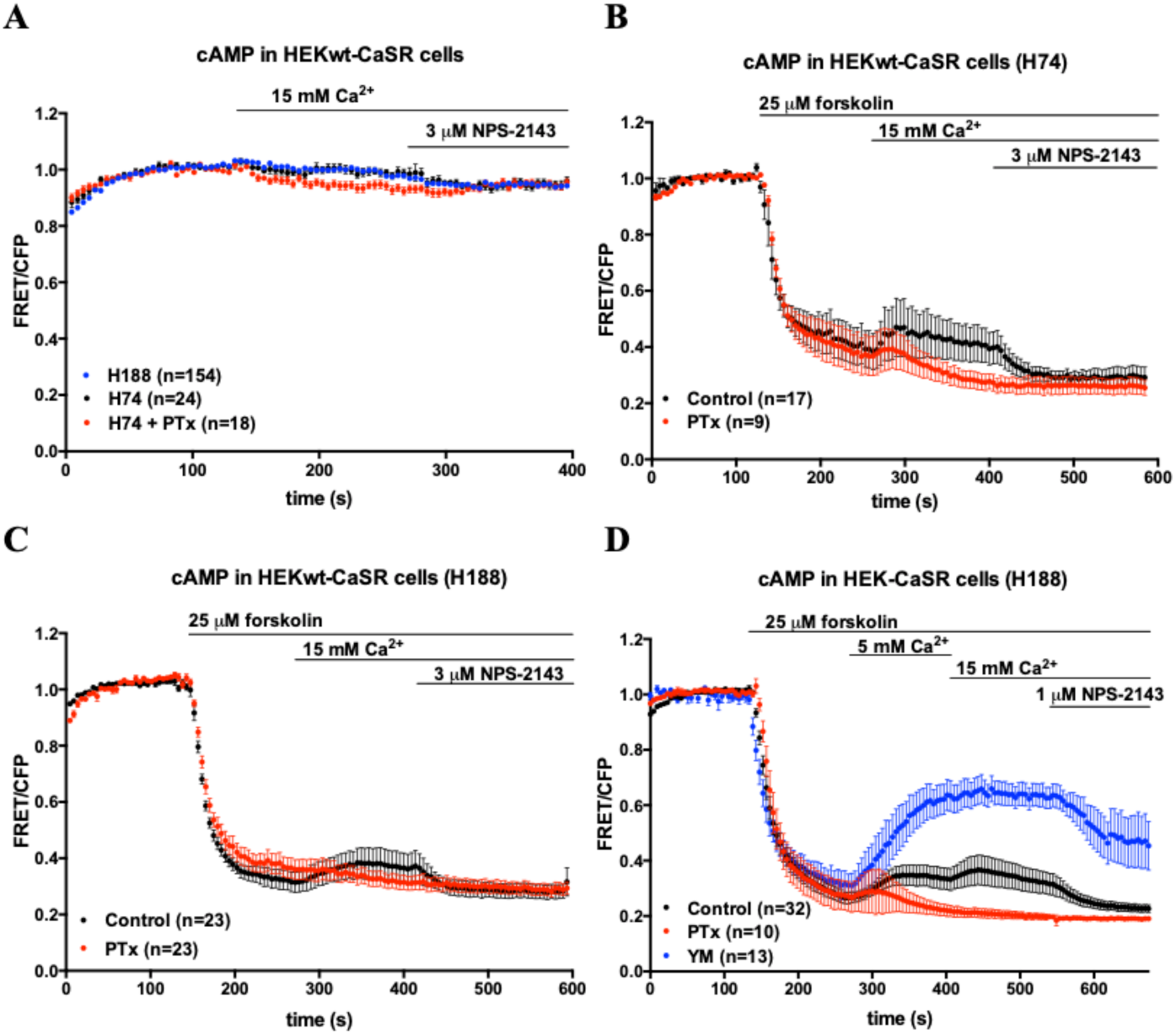
cAMP signaling by CaSR. A: HEKwt cells transfected with CaSR and EPAC sensors H78 or H188. B: HEK-CaSR cells transfected with EPAC sensor H188. Cells were stimulated at the showed time points with forskolin and Ca2+, and the responses antagonized by NPS-2143. Cells were preincubated overnight with PTx or 2h prior to imaging with YM. Time traces show the average ratio change of FRET/CFP fluorescence, and error bars show the 95% CI of the mean. Number of cells is specified for each condition.

### RhoA signaling by CaSR is dependent of Gq

Some studies suggest that CaSR couples and activates G_12/13_ in certain cell lines, including HEK-CaSR [14,15]. We decided to track direct G_13_ activation using a FRET-based biosensor [16] in HEK-CaSR cells. However, we did not observe any changes upon 5-15 mM calcium stimulation and the transfected cells looked very round (data not shown). As a positive control for the G_13_ sensor, we transfected HEK-293 cells with the G13-coupled LPA2 receptor [17] and stimulated the cells with LPA. Surprisingly, we did not observe robust responses either (data not shown). Given the difficulties to use the G_13_ sensor in HEK-293 cells, we decided to use HeLa cells, which are easily transfectable and have been used during the development of the G13 sensor. LPA stimulation of LPA2R transfected cells led to a robust decrease in FRET/CFP ratio, as shown in Figure 7. On the other hand, calcium stimulation in CaSR-transfected cells led to no ratio changes, similarly to histamine stimulation in H1R-transfected cells, which we used as a negative control as H1R does not couple to G_13_.

**Figure 7.**
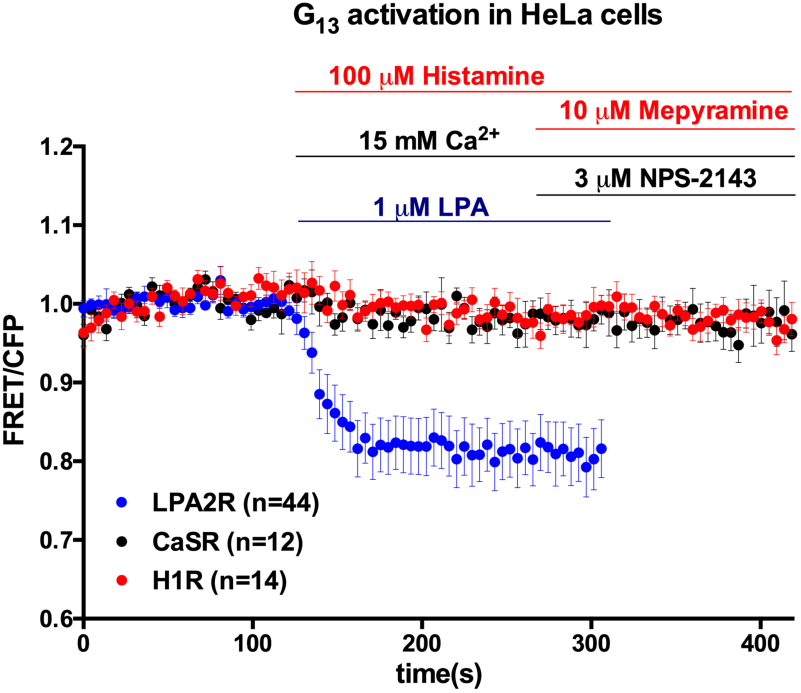
Activation of Gα13 by different GPCRs. Average FRET/CFP ratios plotted over time. HeLa cells were transfected with H1R, CaSR, and LPA2R, stimulated at the showed time points with histamine, Ca^2+^ and LPA, and antagonized by adding Mepyramine and NPS-2143. Time traces show the average ratio change of FRET/CFP fluorescence, and error bars show the 95% CI of the mean. Number of cells is specified for each condition.

Due to cell-type specific differences, it is possible that CaSR does not couple to G_13_ in HeLa cells but it does in HEK-CaSR cells. We then decided to investigate RhoA activity using the DORA-RhoA biosensor [18], as RhoA is a commonly used read-out in G_13_ signaling studies. To identify G_13_-specific signaling we overexpressed the RGS domain of p115RhoGEF to impair G_α13_ signaling [19], as there are no specific chemical inhibitors for G_13_. As shown in Figure 8A, overexpression of this domain abolishes the RhoA response observed upon activation of the endogenously expressed G_13_-coupled protease activated receptors in HEK-293 cells by thrombin. Using HEK-CaSR cells, we observed robust calcium-dependent increases in the RhoA FRET/CFP ratios, which were reversed by NPS-2143. Surprisingly, RGS overexpression did not result in any observable change, as shown in Figure 8B, suggesting that G_α13_ does not mediate CaSR-dependent RhoA activity upon Ca^2+^ stimulation, or that it has a minor role. In addition to G_13_, RhoA activation can also occur downstream of G_q_ proteins, via a pathway that is independent of, and competes with, the PLCβ pathway [20–22]. We found that this was the case in HEK-293 cells transiently transfected with the H1R. Stimulation with histamine resulted in Gq and RhoA activities, that were reversed by adding mepyramine and prevented by inhibition of Gq, as shown in Figure 9. Comparably, preincubation of HEK-CaSR cells with the G_q_ inhibitor abolished the observed RhoA response upon Ca^2+^ stimulation, as shown in Figure 8C. To further confirm this observation, we transfected HEK G_q11_KO cells with CaSR and the RhoA biosensor. As expected, Ca^2+^ addition did not cause any FRET change, and when the expression of Gq/11 proteins was rescued by co-transfecting G_αq_ and/or G_α11_, the RhoA response was restored, as shown in Figure 8D.

**Figure 8.**
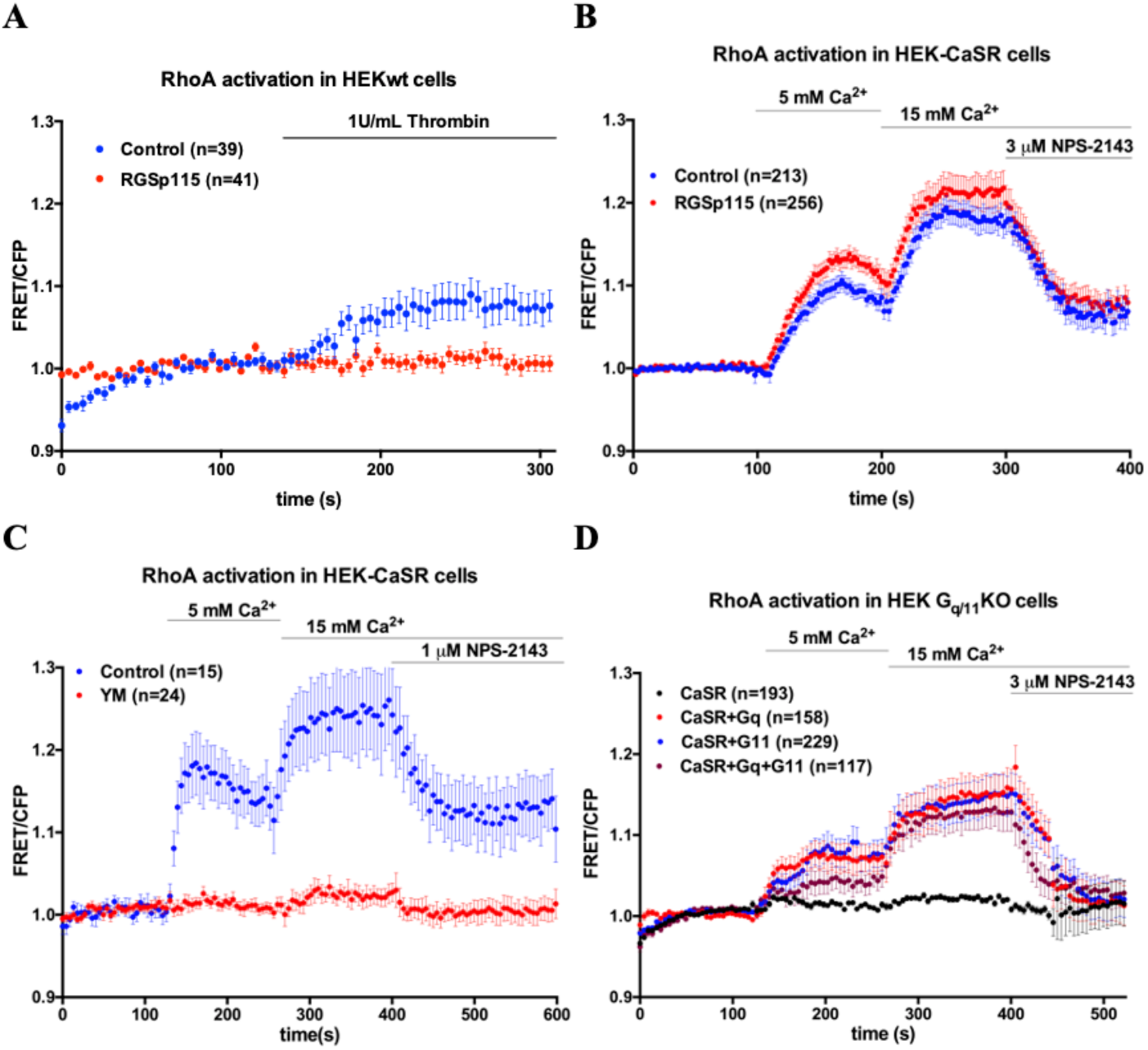
Activation of RhoA by CaSR is dependent on Gq. A: HEK-293 cells were transfected with RGSp115 and stimulated with thrombin to activate endogenous PARs. B: HEK-CaSR cells transfected with RGSp115. C: HEK-CaSR cells in the presence of Gq inhibitor. D: HEK Gq11KO cells were transfected with Gαq and/or Gα11. Cells were stimulated at the showed time points with thrombin or calcium, and CaSR response antagonized by adding NPS-2143. Time traces show the average ratio change of FRET/CFP fluorescence, and error bars show the 95% CI of the mean. Number of cells is specified for each condition

**Figure 9.**
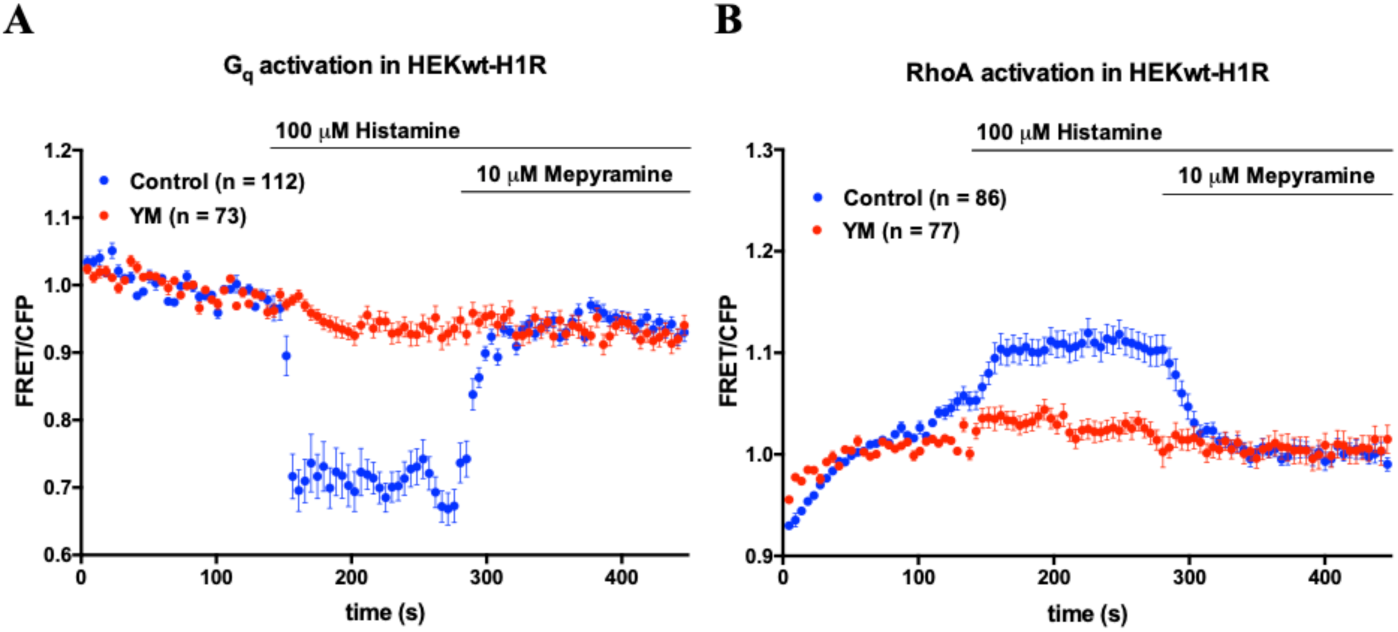
Activation of Gq and RhoA by H1R. A: Average FRET/CFP ratios plotted over time. Cells were stimulated with 100 μM Histamine at the showed time points, and preincubated either with 2 μM YM-254890 or 0.2% DMSO for 2 h prior to imaging. Time traces show the average ratio change of FRET/CFP fluorescence, and error bars show the 95% CI of the mean. Number of cells is specified for each condition.

## DISCUSSION

Signaling immediately downstream of the CaSR has been mainly inferred indirectly and direct activation of the heterotrimeric G proteins has not been observed. Here, we used genetically encoded biosensors that can measure heterotrimeric G-protein activation in single living cells, with high temporal resolution.

Using these biosensors, we observe activation of Gq and the three Gi subtypes upon CaSR activation, in agreement with the well-established coupling of CaSR to the Gq and Gi pathways [6]. The FRET ratios changed rapidly upon stimulation, with comparable kinetics as previously described [8]. While this manuscript was in preparation, another group published similar results using the Gq and Gi2 sensors in HEK-293 cells transiently transfected with CaSR [23]. However, the measured FRET changes kinetics reported by Kienitz *et al*. were considerably slower, with maximum FRET ratio changes observed about one minute after stimulation.

Considering that GPCR and G protein activation typically occur in the millisecond scale, we expect that our observations reflect more accurately the kinetics of these processes.

We have also observed that intracellular calcium mobilization is largely dependent on Gq, and that Gi activity decreases cAMP levels. Single cells exhibited highly diverse intracellular calcium dynamics patterns, from predominantly transient to oscillatory. In sequential stimulation experiments, the calcium responses at a given concentration depended on the previous steps. Further experiments are necessary to understand the mechanisms that modulate the release of intracellular calcium upon Gq activation, and the significance of the different patterns. There are reported cases in literature where calcium oscillations have a signaling advantage over a sustained rise of similar amplitude [24–26], and it is possible too that this is the case for CaSR signaling. By using the single-color intensity-based R-GECO1 sensor, it was possible to simultaneously track Gq activity with the FRET sensor. Furthermore, the clear observation of calcium responses under Gq inhibition conditions could be indicative of residual activity that was not detected by the Gq sensor. This could be explained both by differences in the sensitivity and dynamic range of the sensors and by the characteristic signal amplification of second messengers. Recently, a novel fluorescence lifetime-based calcium sensor was developed [27], based on a circularly permutated fluorescent protein. Tq-Ca-FLITS uses a single CFP channel and provides a quantitative measurement of absolute calcium concentrations, rather than relative changes, simplifying the interpretation of the data and comparison with results from other groups. Alternative variants with other fluorescent proteins will probably be available in the coming years, increasing the versatility of these sensors for multiplexing.

To study cAMP signaling, we tested three variants of a FRET-based EPAC sensor, and found that previous stimulation of the cells with forskolin was necessary to observe any decrease in response to Gi activation by CaSR. To avoid the need of using forskolin to show a decrease in cAMP, the sensors need to be optimized to capture changes at very low cAMP levels, instead of increases [13]. Moreover, we found that the changes in these G protein-preferred downstream events were enhanced when the other G protein was inhibited. It is possible that this results from direct competition between the G protein heterotrimers to bind to the GPCR, or by crosstalk between other mechanisms downstream of the G proteins. It should also be taken into consideration that YM-254890 and Pertussis toxin have different mechanisms of action and binding kinetics. YM-254890 prevents the GDP release from Gαq11 subunits in less than 2 hours [28], while Pertussis toxin needs to be added overnight and catalyzes ADP-ribosylation of the Gαi subunits to prevent coupling to the GPCR [29].

In addition to Gq and Gi pathways, we looked into the G13 pathway, as it has been reported that CaSR can activate G13 in some cell lines [14,15]. The G12/13 family of G proteins is arguably the least characterized of all G protein families, and we attempted to quantify direct G13 activation using the first developed FRET based biosensor for G13 [16]. We observed no effect of calcium stimulation on the single cell FRET ratios, which suggested no detectable G13 activation by the sensor.

Since this sensor had not been used in HEK-293 cells before, we decided to use the G13-coupled LPA2 receptor as a positive control. To our surprise, we did not observe any robust FRET changes after activating the receptor. We also observed that the transfected cells looked considerably rounder, and we believe this occurs due to the overexpression of G13 resulting in increased G13 and RhoA activities. The association between the observed phenotype and high RhoA activity has been described previously in literature [30,31]. Using HeLa cells, on the other hand, we measured robust FRET changes after activating the LPA2 receptor, replicating the results obtained during the engineering and optimization of the sensor. The difference between the results from both cell lines highlights how the dynamic range and usability of a sensor depend largely on the cell line. It is likely that the basal activity of G13 in HEK-293 cells is higher than in HeLa cells and further activation of G13 by the LPA2 receptor escapes the dynamic range of the sensor. Basal G13 activity is affected by factors such as endogenous expression and activity of G13-coupled receptors or regulator of G protein signaling (RGS) proteins, which are unique to each cell type and fluctuate dynamically within single cells.

To explore a possible G13 activation by CaSR in HEK-293 cells, we investigated RhoA activity as it is the main effector downstream of G12/13. Transfection of HEK-CaSR cells with the RhoA sensor, which contains a full-length RhoA, led to a similar phenotype than the observed upon transfection of G13 but we could observe robust RhoA responses upon stimulation with calcium. We found that this activity was abolished by Gq inhibition and not affected by overexpression of the RGS domain of p115RhoGEF, which has GAP activity towards G_α13_. Additional experiments with HEK-293 cells lacking Gq/G11 proteins showed that reconstitution of Gq or G11 restored RhoA activity in response to CaSR activation, supporting the idea that CaSR activates RhoA via Gq and not via G13 in HEK-293 cells. Even though activated RhoA mediates the vast majority of responses downstream of G13 [32], there is evidence of Rho-independent pathways [33–36]. However, to the best of our knowledge, there is no evidence of G13 activation without RhoA-dependent signaling, which suggests that CaSR does not activate G13 as we could only attribute the RhoA response to Gq. Our observations agree with a study that showed that CaSR activates the Gq-RhoA-SRE pathway in HEK-293 cells [37], and are partially supported by a study that found CaSR to promote the assembly of actin stress fibers in a mechanism independent of PTx or PLCβ [38]. Before the development of Gq inhibitors and the accumulation of evidence for Gq-mediated PLCβ-independent mechanisms, PLCβ inhibitors such as U-73122 were used to investigate Gq activation. Moreover, a recent study showed that commonly used concentrations of U-73122 can modulate ion channel activity [39].

Finally, in an attempt to increase the dynamic range of the Gi3 sensor, we replaced cpVenus for mNeonGreen as the acceptor, as it has shown the highest dynamic range in ratiometric FRET imaging using mTq2 as donor in the Gq sensor [7].

However, this change resulted in a significant decrease in the dynamic range, which may be explained by the presence of the Gβ1 subunit in the Gi3 sensor or molecular differences in coupling between the Gαq and Gαi3 subunits.

To summarize, we have shown that Gi and Gq biosensors can be activated by (overexpressed) CaSR and that the activation/deactivation dynamics can be studied in single cells. In combination with inhibitors and biosensors for second messengers, this gives a consistent view of signaling immediately downstream of the GPCR.

### Limitations

In the present study, we used transient transfection to explore the kinetics and activity of multiple signaling molecules using different biosensors. Transient transfection allows to evaluate the effect of genetically encoded constructs in a relatively simple and rapid manner. However, the transfection efficiency is variable between cells, and it is affected by several factors. For instance, upon transfection with the RhoA sensor, we observed a higher fraction of transfected HEK Gq11KO cells than HEK-CaSR cells. In addition, in HEK-CaSR cells, the transfection efficiency with the RhoA sensor was clearly higher than with the different G protein sensors.

Furthermore, the expression levels of the sensors between cells in a single experiment showed variability, which have an effect both on the measured read-outs and in the studied signaling pathways. Similarly, the expression of CaSR can play an important role on the dependent intracellular signaling pathways, as was been shown in literature [40]. By using HEK-CaSR, cell line obtained by clonal selection, we limited the effects of variable CaSR expression due to gene copy number.

We used chemical transfection with PEI due to low cost and simplicity, but the time-consuming generation of stable cell lines would be a valuable effort to obtain large amounts of data and limit the potential consequences of cell-to-cell biosensor expression variability.

Besides the mentioned technical limitations of transient transfection, the overexpression of molecules affects the natural stoichiometry found in the cells and can therefore modify directly or indirectly the behavior of the event of interest. Our observations with the G13 FRET sensor in HEK-293 cells overexpressing the LPA2 receptor point in this direction, however, further experiments are necessary to confirm and explain the apparent unsuitability of this sensor in these cells. For example, using a Tetracycline-dependent system to express the G13 sensor [41] in combination with an additional downstream read-out could show whether the observed morphology and lack of measured FRET changes are due to the overexpression of G13, sensitivity of the sensor, or another factor specific to the LPA2 receptor signaling.

In addition, since the use of G protein FRET-based biosensors involves the overexpression of these proteins, it is possible that heterotrimer dissociation is negligible under normal conditions. In this regard, it is highly recommended to complement the use of these sensors with other (downstream) read-outs to confirm the observations.

## FUNDING

This work was supported by Horizon 2020 Framework Programme (European Union) Award ‘CaSR Biomedicine’, 675228.

## METHODS

### Plasmids

The plasmids C1-Lck-mCherry-RGS-p115RhoGEF and C1-Lck-mCherry were as described previously [19]

The FRET biosensors for Gα_i1,_ Gα_i2,_ Gα_i3_ [8], Gα_q_ [7], RhoA [18], Gα_13_ [16], EPAC [13], and calcium [11] were as described previously. cpVenus was replaced by mNeonGreen in the Gα_i3_ biosensor as previously described [7]. pcDNA3.1 Galpha11 and pcDNA3.1 Galphaq were kind gift of John Sondek. DREADD-hM4Di was obtained from Addgene (plasmid # 45548).

### Reagents

CaCl_2_ dihydrate (Sigma-Aldrich, # 21097) was prepared as a 1M solution in water. Forskolin (Sigma-Aldrich, # F6886) was prepared as a 30 mM solution in ethanol. NPS-2143 hydrochloride (Tocris, # 3626) was prepared as a 1 mM solution in DMSO. YM-254890 (FUJIFILM Wako Pure Chemical Corporation, # 257-00631) was prepared as a 1 μM solution in DMSO. PTX (Invitrogen, # PHZ1174) was prepared as a 100 ng/mL solution in water.

### Cell Culture and Transfection

HEK-293 cells and HeLa cells were obtained from the American Tissue Culture Collection (ATTC, Manassas, VA, USA). HEK-293 Gα_q/11_ KO cells were generated using CRISPR/Cas9 [43] and a kind gift of Dr. Asuka Inoue, Tohoku University. HEK-CaSR cells were a kind gift from Dr. Donald Ward, University of Manchester.

HEK-293 cells, HEK-293 Gα_q/11_ KO cells, HEK-CaSR and HeLa cells were grown in full growth medium, or Dulbecco’s Modified Eagle Medium with GlutaMAX (Gibco, # 61965-059) supplemented with 10% fetal bovine serum (FBS) (Gibco, # 10270106), 100 U/ml Penicillin and 100 μg/ml Streptomycin (Gibco, cat# 15140148).

Two days prior to the experiment, 500 000 - 700 000 cells were seeded on a 35 mm diameter well containing a fibronectin-coated 24 mm diameter glass coverslip (Thermo Scientific Menzel, # 11778691). One day prior to the experiment, the cells were transiently transfected with the constructs of interest using Polyethylenimine (PEI) (1 mg/mL in water). 0.5 μg of DNA per plasmid was mixed with PEI at a ratio 1:4.5 in 100 μL OptiMEM (Gibco, # 31985). When multiple plasmids were transfected, the same ratio and amount of plasmid was used for each of the plasmids.

### Wide-Field Fluorescent Microscopy

The transiently transfected cells were washed 3 times and incubated for 20 min with FBS-free 0.5 mM CaCl_2_ DMEM. The coverslip was then mounted into an Attofluor cell chamber (Invitrogen) in low calcium microscopy medium (20 mM HEPES pH=7.4, 137 mM NaCl, 5.4 mM KCl, 0.5 mM CaCl_2_, 0.8mM MgCl_2_, 20 mM Glucose), and placed in the wide-field fluorescence microscope (Axiovert 200M, Carl Zeiss GmbH) kept at 37 °C, equipped with an oil immersion 40x objective (Plan-Neo-fluor 40× /1.30; Carl Zeiss GmbH). The fluorescence-based biosensors were illuminated using a xenon arc lamp (Cairn Research, Faversham, Kent, UK) with a computer-controlled monochromator. The images were binned 4x4 and recorded with a cooled charged-coupled device camera (Coolsnap HQ, Roper Scientific, Tucson, AZ, USA), controlled with software MetaMorph 6.1 (Molecular Devices).

For FRET experiments, CFP and FRET channels were measured by exciting the sample with 420 nm light (slit width 30 nm), reflecting the light using a 455 DCLP dichroic mirror, and detecting consecutively emission with a 470/30 bandpass filter and 535/30 bandpass filter. Exposure time was 200 ms for CFP and FRET channels and images were acquired every 3-4.6 s. RFP was excited with 570 nm light (slit width 10 nm), and reflected onto the sample by a 585 DCXR dichroic mirror, and emission was detected with a 620/60 bandpass filter.

### Image and data analysis

We used FiJi [44] to extract the fluorescence intensities from the images. First, we manually drew ROIs around the cells, and measured the average intensities in all the channels for each time point in the time-series. To correct for background, we drew an ROI in an area free of cells, and subtracted the average intensity of it to those of the cells. Background-corrected intensities from the FRET channel were corrected for crosstalk from the CFP channel, which was calculated to be 55% as determined by using a CFP-only sample with the same microscope settings. The crosstalk-corrected intensities were then normalised to the first frames, corresponding to those prior to stimulation, and we calculated the ratios of the FRET over the CFP intensities, and plotted them over time using Microsoft Excel.

## Notes

### Competing Interest Statement

The authors have declared no competing interest.

